# Drivers of plant diversity, community composition, functional traits and soil processes along an alpine gradient in the central Chilean Andes

**DOI:** 10.1101/2023.01.13.523936

**Authors:** Lucy Schroeder, Valeria Robles, Paola Jara-Arancio, Cathleen Lapadat, Sarah E. Hobbie, Mary Arroyo-Kalin, Jeannine Cavender-Bares

## Abstract

High alpine regions are threatened but understudied ecosystems that harbor diverse endemic species, making them an important biome in which to test the role of environmental factors in driving functional trait-mediated community assembly processes. In a high mountain system in the central Chilean Andes, we tested hypotheses about the drivers of plant community diversity, functional composition and soil processes along an elevation gradient. We surveyed vegetation and spectroscopic reflectance (400-2400 nm) to quantify taxonomic, phylogenetic, functional, and spectral diversity at five sites from 2400 m to 3500 m elevation. We characterized soil attributes and processes by measuring water content, carbon and nitrogen, and net nitrogen mineralization rates. At high elevation, colder temperatures led to a reduction in available soil nitrogen, while at warmer, lower elevations, soil moisture was lower. Metrics of taxonomic, functional, and spectral alpha diversity peaked at mid-elevations, while phylogenetic species richness was highest at low elevation. Leaf nitrogen followed global patterns of increasing leaf nitrogen with colder temperatures, a pattern consistent at the community level as well as within individual species. The increase in leaf nitrogen, coupled with shifts in taxonomic and functional diversity associated with turnover in lineages, indicate that the ability to acquire and retain nitrogen in colder temperatures may be important in plant community assembly in this range. Such environmental filters have important implications for forecasting shifts in alpine plant communities under a warming climate.

## 1. Introduction

Prevailing theories about community assembly posit that communities are formed from species with functional traits that allow them to disperse into, establish, and persist in the local environment (Bazzaz, 1991; Woodward & Diament, 1991). Trait-mediated assembly processes thus play a critical role in driving the composition and diversity of ecosystems across environmental gradients (Shipley, Vile, & Garnier, 2006). Since Humboldt’s treatment of shifting plant communities with elevation on Mount Chimborazo (von Humboldt, Bonpland, & Montufar, 1824) in the Ecuadorian Andes and his phytogeographic maps of the terrestrial land surface on Earth (Johnston, Brewster, & Berghaus, 1848) scientists have endeavored to characterize changes in diversity across elevational, latitudinal, and other environmental gradients. Deciphering the factors that influence species distributions remains a central issue in ecology and evolution and is paramount to our ability to understand and model how ecological communities are shifting in relation to ongoing climate change. Montane regions of the world harbor unique flora that are highly threatened by climate change (Ackerly et al., 2020; Meng et al., 2019; Wright et al., 2018). The Mediterranean alpine climate of central Chile in particular hosts the highest phylogenetic diversity in the country (Cowling, Rundel, Lamont, Arroyo, & Arianoutsou, 1996; Kalin Arroyo et al., 2006; Scherson, Albornoz, Moreira-Muñoz, & Urbina-Casanova, 2014).

Taxonomic, functional, and phylogenetic dimensions of diversity may have distinct responses to varying environmental conditions along elevation gradients and elucidating these responses can help us understand alpine community assembly (Grime, 2006; Keddy, 1992; Weiher et al., 1998). Often plant species richness peaks at intermediate elevations along montane gradients, which may be due to the alleviation of physiological stress at either end of the gradient which allows for more phenotypes and species to persist (Bryant et al., 2009; Jiang, Ma, & Anand, 2016). Species’ functional traits influence their tolerance to environmental factors and can reflect the ecological pressures that determine species realized assemblages (McGill, Enquist, Weiher, & Westoby, 2006), including along elevation gradients (Fallon & Cavender-Bares, 2018; Read, Moorhead, Swenson, Bailey, & Sanders, 2014; Tang, Morris, Zhang, Shi, & Vesk, 2022). Phylogenetic metrics of diversity provide insight into which lineages can persist, and which species can co-occur locally, across the elevation gradient relative to the available species pool (Cadotte, Albert, & Walker, 2013), which can be useful in inferring community assembly processes (Kraft & Ackerly, 2014). Phylogenetic data along with functional traits can contribute to inferences on whether adaptations to environmental niches are clustered within particular lineages or the extent to which lineages have diversified along environmental gradients (Cavender-Bares, Kozak, Fine, & Kembel, 2009; Tofts & Silvertown, 2000; Webb, 2000).

Spectral diversity integrates chemical, structural, and morphological diversity with light reflectance across a range of wavelengths, providing a high-dimensional, high throughput quantification of plant phenotypes (Kothari & Schweiger, 2022). Functional, phylogenetic, and spectral diversity are closely related in prairie and forest systems (Cavender-Bares et al., 2017a; E. Laliberté, Schweiger, & Legendre, 2019; Schweiger et al., 2018; R. Wang & Gamon, 2019), which is useful for rapidly estimating plant community diversity. We can also use spectra to estimate plant function through the use of spectral indices associated with plant physiology or by using spectrally based models to predict traits from spectral data. However, we do not yet know how spectrally estimated traits are associated with the unusual climate and abiotic conditions of high alpine Mediterranean ecosystems.

In high alpine systems above the tree line, harsh abiotic environmental conditions may be critical in limiting species and lineage distributions. Climatic factors in high alpine Mediterranean ecosystems like high aridity or low temperature can lead to decreased species richness in alpine environments (López-Angulo et al., 2018; Moser et al., 2005). Functional diversity to decreases with elevation in Mediterranean alpine ecosystems, implying that the changes in environmental conditions along the elevation gradient may limit the traits of the plant species there (López-Angulo et al., 2020). In these systems, summer drought and freezing both occur, albeit to different extents along the gradient. Drought impacts water flow in the plant and causes leaf desiccation, meanwhile freezing can damages cells due to ice formation and can also limit water transport by embolisms in the xylem (Cavender-Bares, 2005). However, species with high drought tolerance may have enhanced freezing resistance due to the similar mechanisms by which plants can avoid drought and freezing, such as increased osmolytes, leaves with lower specific leaf area and higher dry matter content, and smaller xylem vessels (Lopez-Iglesias, Villar, & Poorter, 2014; Pescador, Sierra-Almeida, Torres, & Escudero, 2016; Sakai, 1970)

Colder temperatures with increasing elevation limit plant physiological processes, growth and survival, and represent a major selective force and environmental filter (Christian Körner, 2021). Cold temperatures at high elevations require plants to have physiological adaptations to accumulate the necessary resources to maintain their photosynthetic processes at a rate that enables survival. In colder climates the enzymatic reaction rates for photosynthesis are reduced, leading plants to retain higher leaf nitrogen concentrations (Reich & Oleksyn, 2004).

Leaf mass per area is also known to increase under low temperatures both within and across species (Poorter, Niinemets, Poorter, Wright, & Villar, 2009). Limited cell expansion with cold stress can lead to a large number of small cells per unit area and hence to denser leaves (Atkin, Loveys, Atkinson, & Pons, 2006). Denser leaf tissue, in turn, has been shown to reduce freezing damage by slowing the rate of freezing (Ball et al., 2002).

At lower elevations just above the treeline, warm austral summer temperatures lead to high aridity, limiting the water supply to plants. Water is critical to cell expansion and growth as well as fundamental physiological processes such as nutrient transport and photosynthesis (Lambers, Chapin, & Pons, 2008). While low water and low temperature stresses can lead to similar plant physiological and morphological responses, the magnitude of plastic responses can vary within species, and species and lineages adapted to these contrasting stresses are also likely to differ (Poorter et al., 2009). Water-limited and low temperature conditions can reduce photosynthetic rates, preventing photochemical dissipation of absorbed light energy. As a result, plants invest in increased photoprotection through the xanthophyll cycle (Bjorkman, 1987; Demmig-Adams & Adams, 2006; Savage, Cavender-Bares, & Verhoeven, 2009), particularly in the absence of carbon or nitrogen allocation adjustments, such as increased LMA or tissue nutrient concentrations. Photoprotective stress responses can be measured through the photochemical reflectance index (PRI), which indicates the status of the xanthophyll deepoxidation state and total xanthophyll pigment pool size (Wong & Gamon, 2015). A low PRI value can indicate greater deepoxidation and increased plant photoprotection in response to stress.

Temperature can also have an indirect effect on nutrient availability in the soil, which impacts plant growth (Chapin, 1983). The available pool of nitrogen in the soil can impact plant functions and traits, including leaf nitrogen concentration, which is also influenced by leaf mass (Rosati, Day, & DeJong, 2000). Varying trends in soil nitrogen across mountains globally suggest that the context of geography and local plant communities is required to understand local trends in soil nitrogen with elevation. Total soil nitrogen tends to increase with elevation and is affected by vegetation (Decker & Boerner, 2003; Garten Jr., 2004; Smith, Halvorson, & Bolton, 2002), although past studies have largely been conducted in lower altitude systems. Cold temperatures at very high elevations, however, may limit inorganic soil nitrogen availability due to slower microbial nitrogen mineralization rates (Fisher et al., 2013; Körner, Bannister, & Mark, 1986; Nottingham et al., 2015; Pérez-Ramos et al., 2012). In Mediterranean alpine ecosystems, soil nitrogen may impact plant diversity along the elevation gradient by selecting for species that are adapted low nitrogen, resulting in fewer species in sites with lower nitrogen availability and less functional diversity(López-Angulo et al., 2020, 2018).

The purpose of the current study is to advance understanding of the drivers of plant community assembly in threatened high alpine ecosystems. Our study covers the high mountain environment from 2400-3500 m in the Chilean Andes. We use novel and classic methods of assessing plant diversity and composition by incorporating soil chemistry and plant function to discern how abiotic conditions influence plant communities. Along an elevation gradient above the treeline in the central Chilean Andes, we examined how patterns of taxonomic, phylogenetic, functional, and spectral diversity varied among plant communities. Our central goals were 1) to determine how these different dimensions of diversity shift across the elevation gradient, 2) to quantify abiotic environmental conditions across the elevation gradient that might influence plant community assembly, and 3) to characterize relationships between plant traits and changing abiotic conditions along the elevation gradient to unveil potential abiotic drivers of community assembly.

Following our first goal, we hypothesized that changes in aridity and temperature along the elevation gradient would drive shifts in plant composition that influence diversity patterns. Temperature can affect other environmental conditions, which in turn have influence which plants can survive in those conditions. For our second goal, we then hypothesized that i) soil water would increase with increasing elevation where air temperatures and evaporative demand were lower, and ii) soil microbial activity would decline with decreasing temperature at high elevations, such that net nitrogen mineralization rates would decline with increasing altitude resulting in less available soil nitrogen. Such changes in environment can select for plants with certain traits to survive such conditions, affecting diversity. For our final goal, we hypothesized that shifts in plant traits both within species and communities across the gradient would correlate with i) increasing cold stress with increasing elevation. As such, we predicted higher LMA, chlorophyll a concentration, and leaf nitrogen concentration at the coldest sites, and ii) higher aridity and greater water stress at the lower end of the gradient resulting in lower leaf water content and lower PRI. We anticipated that both cold and aridity stress would be low in the middle of the gradient, resulting in the most diverse plant communities.

## 2. Methods

### 2.1 Field Site

Our field sites were in the highly diverse Región Metropolitana in the Chilean Andes (33°S, 70°W). Shrubs and annual plants are restricted to lower elevations, whereas cushion plants and perennial rosettes dominate in mid to high elevations above the treeline (Cavieres, Peñaloza, & Kalin Arroyo, 2000). We sampled at five sites in this area along an elevation gradient from 2400-3500 m (supplemental figure 1). This elevation gradient encompassed a temperature gradient with a minimum January (summer) temperature of 2.1 °C at the highest elevation, and 7.8 °C at the low end (Cavieres, Badano, Sierra-Almeida, & Molina-Montenegro, 2007).

At each of the five elevations we established 10-meter transects and sampled the plant diversity using a point-intercept method to document every plant within a 10 cm radius at every 0.5 meter point along the transect. Each site had seven transects, except the highest elevation site which had eight transects to better capture the total diversity despite extreme sparseness. At this site, every plant within a 10 cm radius of the transect was sampled to detect rare species. In plant cover calculations, to ensure cover at this site was not over estimated using this method compared to the other sites surveyed every 0.5, we only included plants occurring at every 0.5 meters along the transect.

### 2.2 Spectral data collection

We collected hyperspectral data for every plant sampled along the transect (every 0.5 m) using a SVC HR 1024i spectroradiometer and leaf clip foreoptic. We removed the leaves from each plant and measured spectral reflectance within two hours. For plants with small or narrow leaves, we used more than one leaf from the same individual without overlapping to fully cover the foreoptic. A white reference was taken every ten measurements to calibrate the instrument. The spectra were compiled and trimmed to only include wavelengths from 400 to 2400 nm to remove noisy sections at the edges of the sensor’s detection range RStudio version 4.1.2 (R core team, 2021) using the R package spectrolab version: 0.0.16 (Meireles, Schweiger, & Cavender-Bares, 2021).

### 2.3 Canopy data collection

We chose a single site (2800 m) with high vegetation cover to collect canopy-level spectral data. In each of the 10 m transects, we created 40 0.25 × 0.25 m quadrats. We measured hyperspectral reflectance from 1 m above each quadrat with 4° foreoptic lens. In addition, we categorized the percent vegetation cover in each quadrat. In every fourth quadrat, we collected a 5 cm diameter soil core to 10 cm depth.

### 2.4 Soil measurements

We sampled soil cores to 10 cm depth along the elevation gradient to determine nutrient variation and the soil nutrient availability. At each site, we took five samples on bare soil near the transects and five samples under the species *Phacelia secunda*, which is present across the elevation gradient. Samples under *P. secunda* allowed us to directly correlate the plant function with the soil conditions below it. The bare soil samples allowed us to estimate the soil conditions without effects of the plants growing above the sampled area. The soil was sieved with a 2 mm mesh sieve to remove large rocks and organic matter. We measured the wet and oven-dry weight of the soil samples to determine gravimetric water content. To measure the total soil carbon and nitrogen concentration, the dried soil was ground to a homogenous fine powder and analyzed with a Costech ECS4010 elemental analyzer (Valencia, California, USA). We measured pH in water by mixing 10 g of the oven-dry soil with 20 mL of nanopure water and measuring the pH of the extract with an Orion benchtop model 420A pH probe (Beverly, MA, USA).). Fresh soils were shipped on ice to the University of Minnesota where we performed an initial extraction in KCl, established room-temperature laboratory incubations in the dark within 12 days of field collection to measure net nitrogen mineralization over 30 days. Net nitrogen mineralization rate Net nitrogen mineralization was calculated as the difference between initial and final extractable concentrations of ammonium and nitrate + nitrite-nitrogen, which were measured using methods adapted from Doane and Horwáth (2003) (see supplemental methods for additional detail).

### 2.5 Phylogeny construction

#### 2.5.1 Taxon sampling

DNA was obtained from leaf material of individuals collected in the field and from herbarium material stored at CONC (Herbarium of the Department Botany, University of Concepcion) and SGO (Herbarium of the National Museum of Natural History). Samples were stored in silica gel. Vouchers for field-collected material are deposited in the herbaria CONC. We downloaded sequences for some species from GenBank (NCBI; Supplement Table 1).The total number of taxa considered was 64. *Ginko biloba* (Ginkgoaceae) was chosen as an outgroup.

#### 2.5.2 DNA extraction, amplification and sequencing

Genomic DNA was extracted with the DNeasy Plant Kit (Qiagen, Valencia, CA, USA). We amplified the DNA with PCR using the primers listed in supplemental table 1 (see supplemental methods for additional detail). Samples were sent to Macrogen (Seoul, South Korea) for purification and sequencing. Sequences were loaded, edited and aligned using ChromasPro 2.33 (Technelysium, Brisbane, Australia) and BioEdit 7.0 (Hall, 1999) and have been deposited in GenBank (Table 1).We performed a combined analysis for sequences of the nuclear gene ITS and two chloroplast genes, *rbc*L and *matK*. Bayesian inference analyses were performed with MrBayes (Ronquist et al., 2012) (see supplementary methods for details of Bayesian analysis).

**Table 1.** Spectral indices used to acquire phenotypic data from leaf level hyperspectral reflectance data.

| Index | Equation | Description | Citation |
| --- | --- | --- | --- |
| NDMI | $(R_{1649} - R_{1722}) / (R_{1649} + R_{1722})$ | Normalized Dry Matter Index | (L. Wang, Hunt, Qu, Hao, & Daughtry, 2011) |
| LMA | $(R_{860}/R_{1240})$ | Simple Ratio Leaf Mass per Area | (Q. Zhang, Li, & Zhang, 2012) |
| LWP | $(R_{1200}/R_{850})$ | Simple Ratio Leaf Water Potential | (Fujino, Endo, & Omasa, 2002) |
| RWC | $(R_{900}/R_{970})$ | Simple Ratio Relative Water Content | (Penuelas & Inoue, 1999) |
| PRI | $(R_{531} - R_{570})/(R_{531} + R_{570})$ | Photochemical Reflectance Index | (Gamon, Serrano, & Surfus, 1997) |
| ChlNDI | $(R_{750} - R_{705})/(R_{750} + R_{705})$ | Chlorophyll A normalized difference index | (Gitelson & Merzlyak, 1994) |

### 2.6 Spectral indices

We used reflectance bands to calculate six spectral indices related to photosynthetic function, water capacity, and leaf structure.

### 2.7 Canopy NDVI

We calculated the normalized difference vegetation index (NDVI) using the equation NDVI=(R_900_-R_685_)/(R_900_+R_685_) following methods by Penuelas and Inoue (1999). We compared the mean NDVI and mean vegetation cover for each transect and compared NDVI with soil nitrogen for each quadrat at 2800 m.

### 2.8 Leaf tissue traits

Leaf tissue was collected and dried after taking a spectral measurement. We randomly selected 233 samples, making sure to include samples of each species at each elevation. The dried leaf tissue was ground with a Wiley mill using a 40 mesh (0.42 mm) screen, then analyzed with Costech ECS4010 elemental analyzer (Valencia, California, USA) to determine carbon and nitrogen concentrations.

Percent leaf nitrogen for all individuals was predicted using spectral measurements by fitting a partial least squares regression (PLSR) model following methods described by Schweiger et al (2018). Our percent nitrogen dataset included measurements from 233 individuals, each matched with spectral data. We used 80% of the data to create a model over 1000 iterations, using the plsr() function from the R package pls version 2.8.0 (Hovde Liland, Mevik, Wehrens, & Hiemstra, 2021) and validated the model on the remaining 20%, resulting in a RA^2^ of 0.42 and a standardized RMSEP of 0.48, indicating that the traits were predicted moderately well. We extracted the coefficients from this model and applied them to the remaining spectral data to calculate the percent nitrogen for all the individuals (n=836) in our dataset. For consistency, we used spectrally estimated percent leaf nitrogen for all individuals in our analyses.

### 2.9 Multiple measures of alpha diversity

#### 2.9.1 Taxonomic diversity

To compare the number of species at each site, we calculated species richness and species evenness at each transect. We calculated Shannon diversity index in the R software package vegan version 2.5.7 (Oksanen et al., 2019) and divided it by the natural log of the number of species in each site to measure species evenness using the Shannon equitability index (Shannon & Weaver, 1949).

#### 2.9.2 Phylogenetic diversity

We made the phylogeny ultrametric using the nnls method with the force.ultrametric() function of phytools version 1.0.1 (Revell, 2012). Using the phylogeny and species abundance dataset, we calculated several metrics of phylogenetic diversity to compare the relatedness of species at each site. Using functions in the package picante version 1.8.2 (Kembel et al., 2010), we calculated phylogenetic species richness (PSR) which is comparable to species richness, but accounts for phylogenetic relatedness, and phylogenetic species evenness (PSE), which incorporates species abundances with phylogenetic relatedness and richness (Helmus, Bland, Williams, & Ives, 2007).

#### 2.9.3 Functional diversity

Seven traits including spectrally predicted leaf nitrogen and the six spectral indices (table 1) were centered and scaled using the base R version 4.1.2 function scale() for the functional diversity calculations (Becker, Chambers, & Wilks, 1998). We then created a Euclidean distance matrix from these measures. Using this distance matrix and a species abundance matrix, we calculated metrics of functional diversity of each transect using the dbFD function from the FD package version 1.0.12 (Laliberté, Legendre, Shipley, & Laliberté, 2015). This function uses a distance-based framework to calculate multidimensional functional diversity indices including functional evenness and functional dispersion, which respectively measure the homogeneity of functional traits and the distance from the centroid of functional traits.

#### 2.9.4 Spectral diversity

We calculated spectral diversity to incorporate all the information that the full spectrum encompasses. We used the same Euclidean distance-based method as described for the functional diversity measure, except that we used spectral reflectance values for each wavelength band as the input for trait values, following methods used by Schweiger et al (2018).

We regressed metrics of taxonomic, phylogenetic, functional, and spectral diversity against elevation to understand the patterns of alpha diversity along our elevation gradient. To decide if the best fit was linear or polynomial, Akaike information criterion (AIC) values were calculated and considered with the overall pattern of the data.

### 2.10 Community composition

We calculated taxonomic, phylogenetic, functional, and spectral composition of communities using NMDS ordinations of distance matrices. For taxonomic composition, we calculated the fractional cover of each species within each of the 10 m transects by counting the abundance of each species in a transect, with points without a species counting as zero. The count of each species was divided by 20, for the 20 points along the 10 m transect. We calculated Euclidean distances between transects based on the abundances of each species to create an abundance-weighted community distance matrix. For phylogenetic composition, we created abundance-weighted community matrices for phylogenetic composition using phylogenetic mean pairwise distances between taxa in each transect using the comdist() function in the R package picante (Kembel, 2010). We calculated functional composition with community-weighted trait Euclidean distances between transects using the function dist() in the R stats package (Becker et al., 1998; Borg & Groenen, 1997; Mardia, Kent, & Bibby, 1979). Similarly, we calculated spectral composition by treating each band of the spectra as a trait and calculated a Euclidean distance matrix with community-weighted means of each transect. NMDS ordinations were conducted on each of these matrices using the metaMDS() function in the R package vegan (Oksanen et al., 2019). We also fitted the environmental variables to the ordination using the envfit() function in the R package vegan to determine if communities clustered along environmental gradients (Oksanen et al., 2019). The first NMDS axis was extracted and plotted against elevation to show the change in community taxonomic, phylogenetic, and functional diversity across the elevation gradient.

### 2.11 Trait-environment associations

We examined patterns of traits, soil nitrogen, and soil water content at the community and species level over elevation. We chose to run these models with a reduced number of traits from table 1 that would inform us on specific aspects of plant function. The traits we selected included percent leaf nitrogen, PRI, Chl a NDI, LMA, and RWC, which informs us on leaf nutrient levels, stress, photosynthetic capacity, leaf economic spectrum, and water status, respectively. For the community level, traits were aggregated by species. We used phylogenetic generalized least squares (PGLS) regression to examine patterns of traits with environmental variables while accounting for species relatedness. We selected *Phacelia secunda* to analyze intraspecific trends in traits with environment. For this analysis, we used regressions of each individual’s trait with environmental conditions. AIC values and visual fit of the points were used to determine linear versus non-linear fits for the model. We used the Holm-Bonferroni step down method to correct our p values from and reduce false positives (Holm, 1979). We chose this method as it is the second-most conservative p value correction method, while being uniformly as powerful as the most conservative method, the Bonferroni method, and would allow for interpretation of the results but avoid inaccurate conclusions based false positives.

### 2.12 Phylogenetic signal

Blomberg’s K (Blomberg, Garland, & Ives, 2003) was used to compare phylogenetic signal in spectral indices. We tested for phylogenetic trait conservativism using a white noise model and a Brownian motion model following Cavender-Bares and Reich (2012) Olalla-Tarraga et al. (2017), Fontes et al. (2022). The white noise model was constructed by randomizing tips across the phylogeny over 1000 simulations and testing how the true K differed from the distribution produced by the randomization procedure. Similarly the Brownian motion model was modeled as a “random walk” of trait evolution over 1000 simulations, bounded by positive and negative infinity. Calculating the statistic under these two models allowed us to interpret the degree of trait conservativism. Significant p-values (<0.05) in the white noise test indicate that the traits are more conserved than they would be by random chance. If the p-value is significant under Brownian motion, it indicates that the trait is not consistent with a Brownian motion model.

## 3. Results

### 3.1 Environmental variation with elevation

Soil water increased linearly as elevation increased (Fig.1a). Net nitrogen mineralization rates in short-term laboratory incubations (measured at a single, common temperature) declined with increasing elevation (Fig.1b). decreased with increased elevation (Fig.1c), as did carbon (not shown). There was no change in net nitrogen mineralization per g of soil nitrogen with elevation or of net nitrogen mineralization with the soil carbon:nitrogen ratio, indicating that the observed change in net nitrogen mineralization is driven by a decline in the quantity of organic soil nitrogen available to be mineralized, rather than the quality of soil nitrogen.

**Figure 1.**
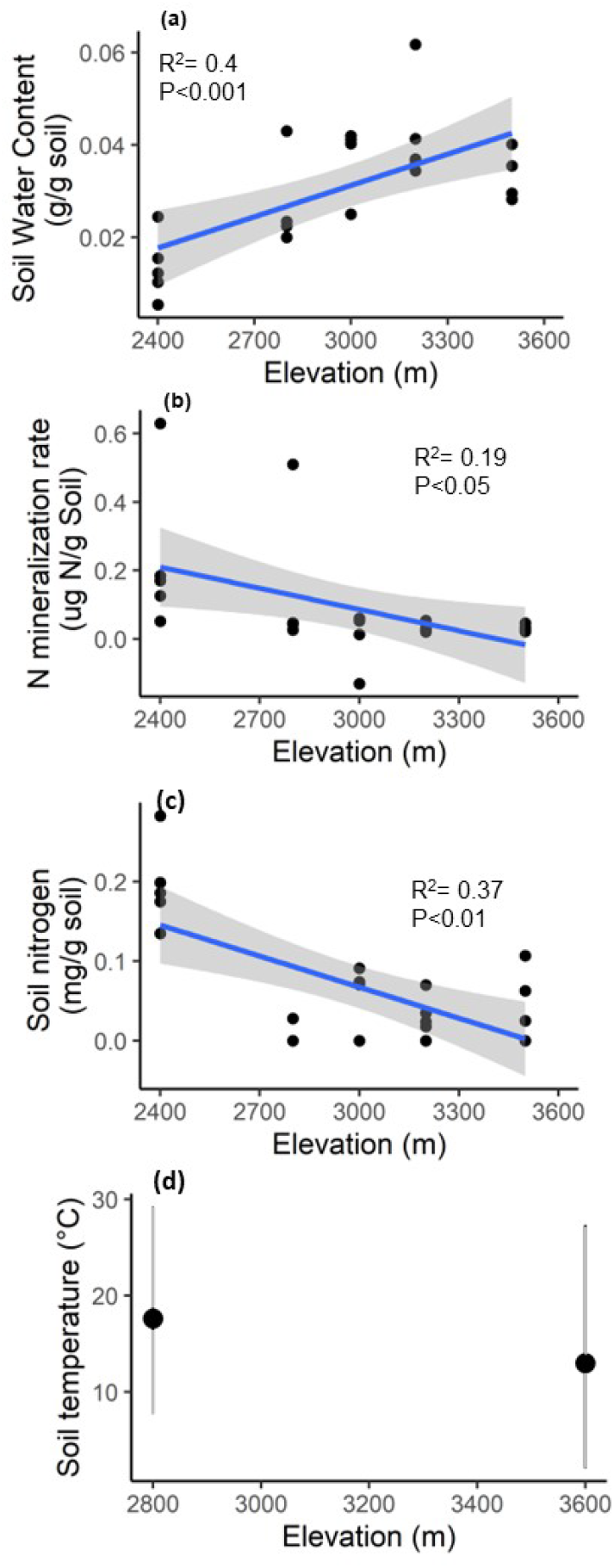
Abiotic and biotic soil variation with elevation. (a) gravimetric soil water content (b) rate of microbial net nitrogen mineralization in 30 laboratory incubations at room temperature (c) bulk nitrogen present in the soil (d) January soil temperature from Cavieres et al (2007), with vertical gray lines indicating daily range and black circles indicating daily mean temperature. Each point represents a transect sample. The best fit line is plotted in blue, with gray shading indicating standard error.

### 3.2 Changes in diversity indices with elevation

We observed mid-gradient peaks in most measures of diversity, including species richness, functional dispersion (FDis), functional evenness (FEve), and spectral dispersion (spec dis) (Fig. 2a,c,g,d). Species evenness (Shannon equitability index) decreased with increasing elevation.

**Figure 2.**
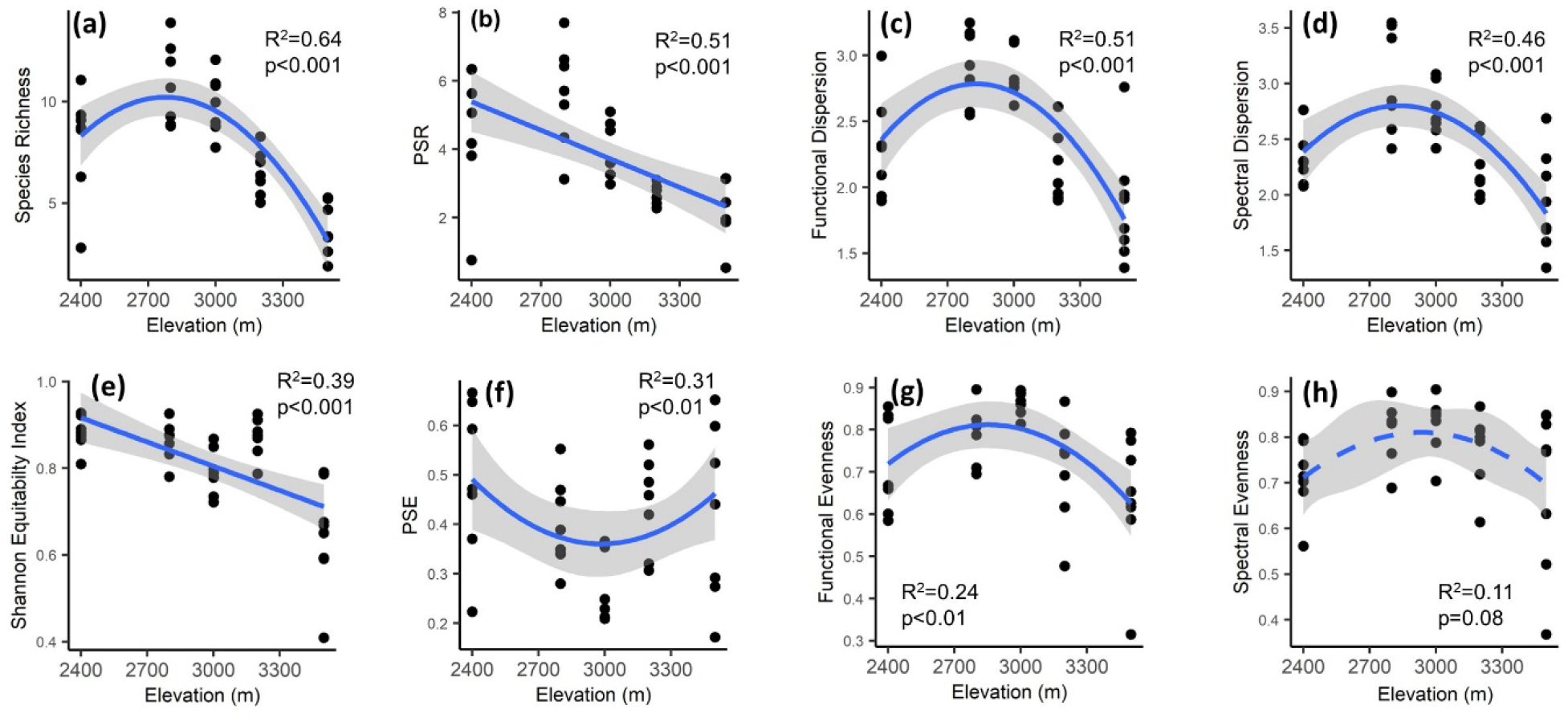
Alpha diversity across an elevation gradient. Taxonomic diversity (a) indicated by species richness indicates that the highest number of species occurred at the lowest elevations, peaking at 2800 m Species evenness (Shannon equitability index) (e) decreased with elevation. Phylogenetic diversity in terms of phylogenetic species richness (PSR) (b) and phylogenetic species evenness (PSE) (f) indicates that as elevation increased, there were fewer species that were less related to each other. Functional dispersion (c) indicates that trait values became more similar at low and high elevations, with a peak in the middle elevation 3000 m. Similarly, spectral dispersion (d) indicates that the hyperspectral signatures were less similar at middle elevations and most similar at the highest elevation. Functional evenness (g) and spectral evenness (h) followed the same trend as dispersion, however the trend was not statistically significant for spectral evenness.

Phylogenetic species richness (PSR) decreased linearly with elevation, whereas phylogenetic species evenness (PSE) was lowest at the sites at the middle of our elevation gradient (Fig. 2b,f). With a p value of 0.08, there was weak evidence that spectral evenness (spec eve) peaked at middle elevations.

### 3.3 Testing for phylogenetic signal in species altitudinal distributions

The range distribution plot (Fig. 3) shows that many species occurred at multiple elevations with shrubs distributed only in lower to mid elevations. There was no obvious phylogenetic pattern in how species were distributed across the elevation gradient. Using Blomberg’s K to analyze phylogenetic signal in traits of all species in the study, we found significant (p<0.05) non-random signal in spectral indices corresponding to relative water content (RWC), leaf mass per area (LMA), and leaf water potential (LWP), and leaf nitrogen using a white noise null model. Under a Brownian motion null model, the observed K values are statistically significant (p<0.05) (see supplemental table 2).

**Figure 3.**
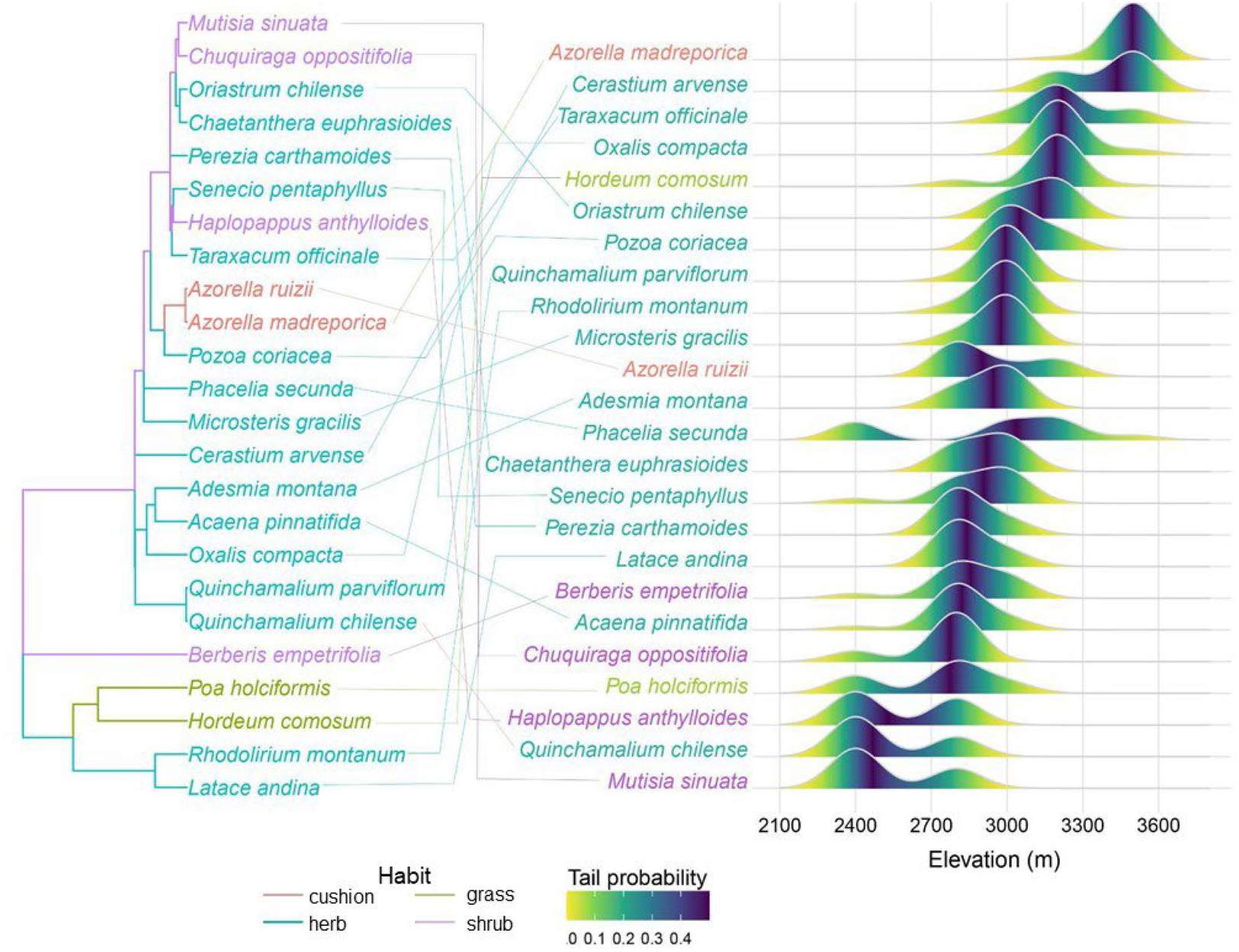
Elevation distribution of species occurring at multiple sites alongside the phylogeny. The ridgeplot indicates the density of each species at each elevation. Species names are color-coded to indicate plant habit. The color gradient in the ridges indicates the tail probability, with the darkest purple color indicating highest density and yellow indicating lowest density.

### 3.4 Shifts in community composition with elevation

Species community composition clustered by site. The increase of species community composition with elevation indicates closer sites had more shared species (Fig. 4a,e).

**Figure 4.**
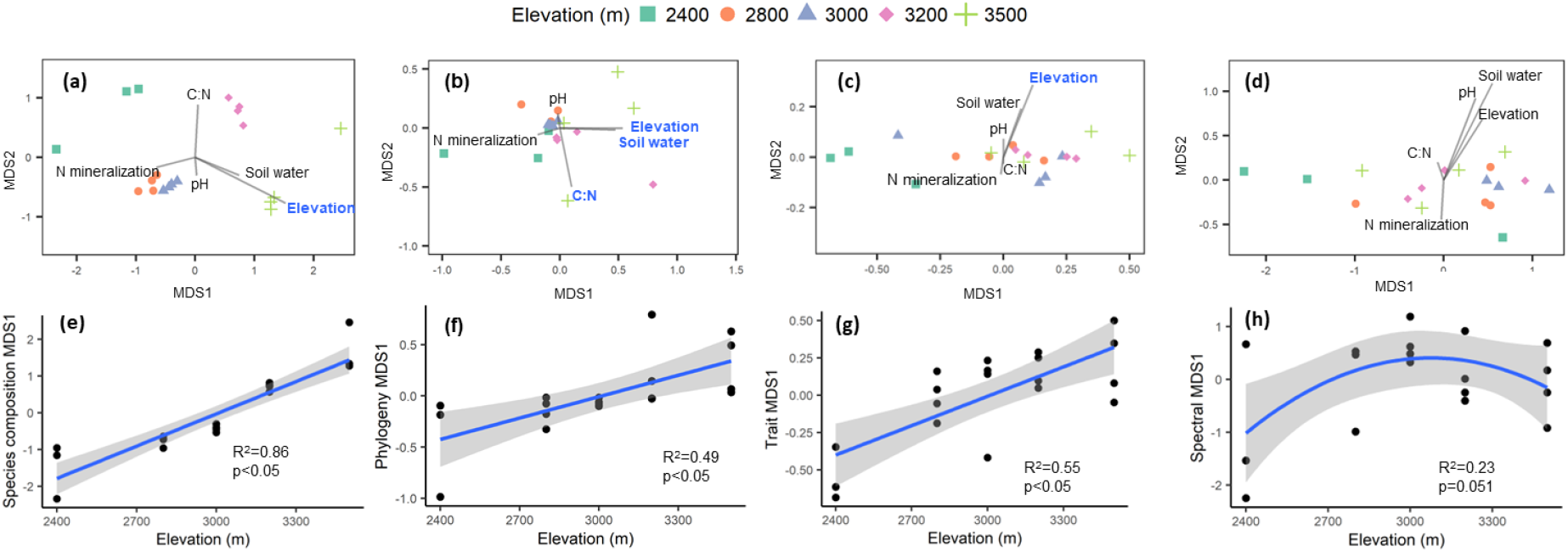
Turnover in composition with environmental variables across the elevation gradient using NMDS ordination. Significant (p<0.05) vectors resulting from the envfit are highlighted in bold blue text. (a) Species community composition calculated as cover per species clustered by site and increased linearly with elevation (e). (b) Phylogenetic community calculated as branch distances did not cluster as clearly but had a similar linear trend with elevation (f). (c) Functional trait composition calculated as trait distances showed traits were mostly distributed along the first MDS axis. (g) Changes in trait composition increased linearly with elevation. (d) Spectral composition did not distinctly cluster, peaking at mid elevations (h).

**Figure 5.**
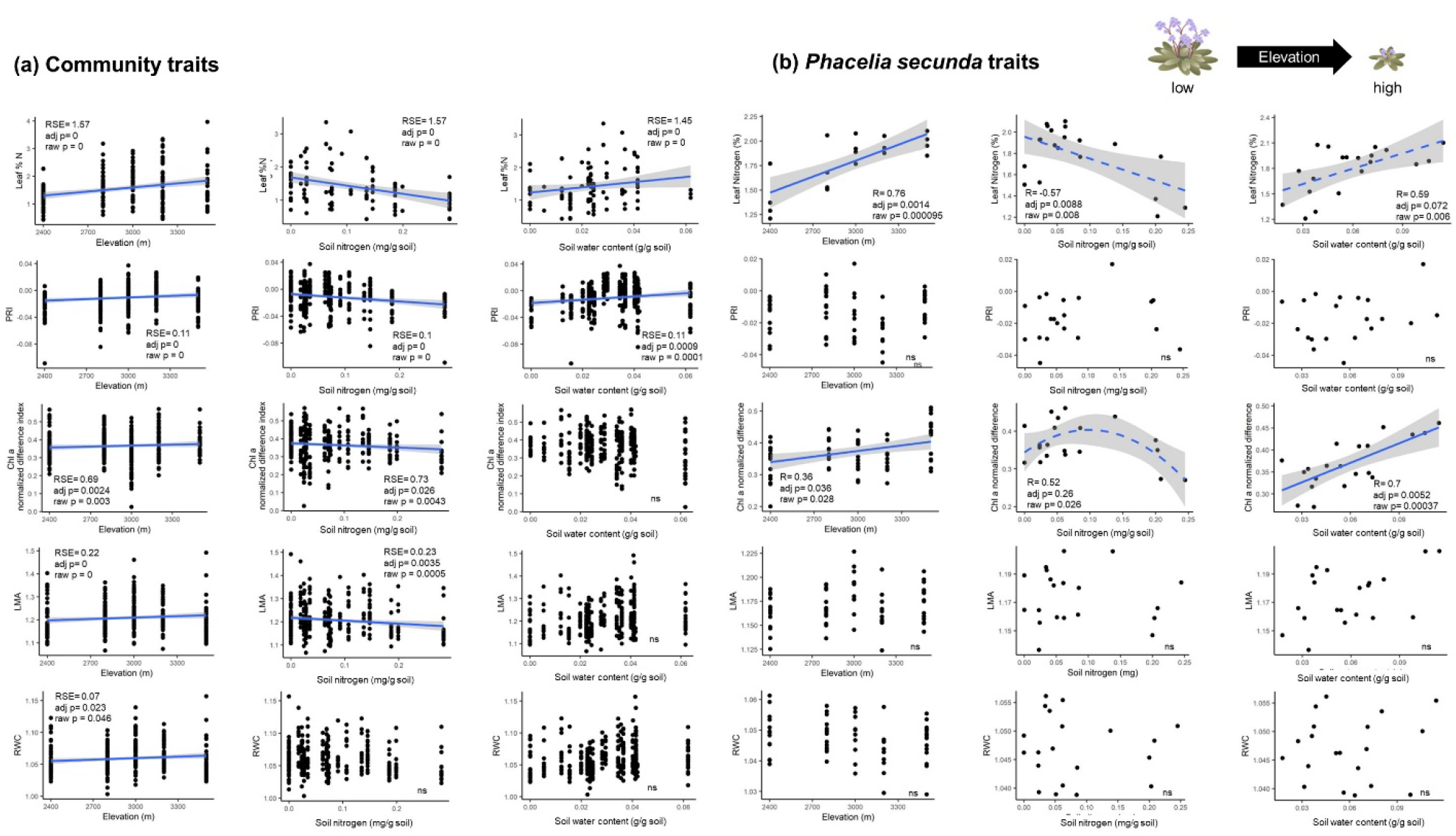
Trends in leaf traits calculated from spectra with elevation and soil conditions of nitrogen and water content at the community level (a) and (b) at the intraspecific level in *P. secunda*, a species found across the elevation gradient. As pictured by the cartoon, the species form transitioned from a tall, shrub-like habit at lower elevations to a small rosette at higher elevations. Solid lines indicate significant relationships with p values adjusted using Holm-Bonferroni step-down correction (*α*=0.05), and p values significant at *α*=0.1 are indicated by dashed lines. Non significant relationships are indicated by “ns”

Phylogenetic composition did not cluster obviously by site but increased linearly when NMDS axis 1 was plotted against elevation, suggesting that species at opposite ends of the gradient were more distantly related (Fig. 4b,f). The first phylogenetic axis also had a significant correlation with soil water and soil C:N. Community composition of traits was dispersed mostly along the first MDS axis (Fig. 4c). The communities at the lowest elevation did not share characteristics with the communities at the highest elevations. The first trait axis the differences in elevation (Fig. 4g). Spectral composition was also dispersed along the first MDS axis, but showed a unimodal relationship with elevation (Fig. 4d,h).

### 3.5 Trait variation among communities and within species across environment gradient

Leaf nitrogen increased with increasing elevation, decreased with increasing soil nitrogen, and increased with increasing soil water. PRI, the index of photosynthetic stress, increased with increasing elevation and soil water, and decreased with increasing total soil nitrogen. The normalized difference index for chlorophyll a (Chla NDI) shows chlorophyll a concentration significantly increasing with elevation, but decreasing with soil nitrogen. The structural index, LMA show that leaves were more dense or thick at higher elevations but decreased with higher soil nitrogen. The index relating to water status, RWC, suggests that plants had more water in their leaves with increasing elevation. We also compared spectral measurements of leaf canopy cover (NDVI) to plant cover and soil nitrogen concentration. NDVI was positively correlated with plant cover and with soil nitrogen (see supplemental figure 3).

Leaf nitrogen increased with elevation in *P. secunda*, following the community-level trend. Leaf percent nitrogen also showed a marginally significant decrease with increasing soil nitrogen and increase with soil water content. The Chl a NDI shows chlorophyll a concentration increasing with elevation and with increasing soil water content. While the corrected p value was not significant, the relationship of chlorophyll a and soil nitrogen was well fit when modeled as a second order polynomial peaking at intermediate nitrogen concentrations and dropping off at high soil nitrogen concentrations.

## 4. Discussion

Along an elevation gradient in alpine ecosystems in the Chilean Andes, shifting environmental factors, including microbially-mediated soil mineralization processes that are coupled with climate, appeared to drive patterns of plant functional, phylogenetic, and spectral diversity. Soil nitrogen availability declined with elevation, yet spectrally determined leaf nitrogen concentration increased both within individual species and in whole communities. This pattern is consistent with global patterns of increased leaf nitrogen accumulation in low temperatures to compensate for slower photosynthetic reaction rates (Körner, 2021; Oleksyn et al., 1998; Reich & Oleksyn, 2004; Woods et al., 2003). We found higher taxonomic, functional, and spectral diversity at middle elevations, consistent with our hypotheses and previous work (Bryant et al., 2009; Jiang et al., 2016). However phylogenetic diversity differed from this trend: some lineages occurred only at lower elevation sites, thus phylogenetic species richness was highest just above the tree line and decreased with elevation. Species that are adapted to the contrasting conditions at either end of the elevation gradient lead to changes in community composition. The combined patterns of plant diversity and composition indicate that environmental filters likely drive plant community assembly along the elevation gradient and that the nature of these filters shift with altitude, similar to other montane systems (Chun & Lee, 2018; Fallon & Cavender-Bares, 2018; Scherrer et al., 2019; Yi et al., 2020; Zhang, Huang, Wang, Liu, & Du, 2016)(Chun & Lee, 2018; Fallon & Cavender-Bares, 2018; Scherrer et al., 2019; Yi et al., 2020; W. Zhang et al., 2016).

### 4.1 Environmental conditions influence physiological traits along the elevation gradient

Harsh environmental conditions at both ends of the gradient select for species with specific adaptations that enhance survival under those specific conditions. The low end of the elevation gradient was characterized by lower soil water availability, warmer temperatures, and more nitrogen availability, selecting for plants that can survive with low leaf water content.

However the trend in water content may also be explained by the thinner leaves at low elevations compared to high elevations, with high elevation leaves having both more dry matter and more water due to slower growth rates at low temperatures (Berry & Bjorkman, 1980). Lower net nitrogen mineralization rates measured at the highest elevation suggest that temperatures reinforce lower nutrient availability in soils measured from that site, amplifying the reduction in nitrogen availability with increasing elevation (Guntiñas, Leirós, Trasar-Cepeda, & Gil-Sotres, 2012). These reduced net nitrogen mineralization rates are driven by a reduction in the amount of organic soil nitrogen with increasing elevation, as evidenced by the lack of effect of elevation on nitrogen mineralization per gram of soil nitrogen. Lower plant cover at this higher elevation likely impacts the amount of organic nitrogen inputs into the soil. The positive relationship of canopy NDVI with soil nitrogen at the 2800 m mid-elevation site supports the importance of soil nitrogen availability for plant growth.

Decreased nutrient availability due to colder temperatures at the higher elevations likely drives differences in plant communities along the gradient by selecting for species with small, dense leaves (as indicated by the increased LMA) and high nitrogen accumulation ability, despite low soil nitrogen. Increased leaf nitrogen and higher LMA may be linked, resulting from increased thickening of the cell walls (Onoda et al., 2017). However, the higher photosynthetic chlorophyll a pigment concentrations at higher elevations suggests that the leaf nitrogen accumulation at these elevations may play a role in the maintenance of photosynthetic activity Leaf nitrogen concentration is correlated with the concentrations of chlorophyll, which, and RuBisCO (Ellis, 1979; Evans, 1989; Luo et al., 2021). High RuBisCO concentrations help to maintain photosynthetic efficiency in cooler temperatures where biochemical reaction rates are slower (Cerqueira et al., 2019; Holaday et al., 1992).

### 4.2 Intraspecific trait variation allows some species to survive both harsh environments

Species that survive across the gradient may be able to respond plastically to or are locally adapted to different environmental conditions. Trends in leaf nitrogen for *Phacelia secunda*, which occurs across the gradient, were consistent with variation across communities, suggesting intraspecific variation in a trait that is important for survival at different elevations. *P. secunda* had high dry matter content in both the lowest elevation and highest elevation. High dry matter content in alpine plants helps plants maintain turgor at lower water potential, which may help *P. secunda* to survive both drought and cold stress (Pescador et al., 2016). Such trait plasticity may be important to facilitate plant migration and survival in mountains, especially under climate change (Henn et al., 2018). The trait plasticity in species like *P. secunda* may also allow some species that are highly abundant at intermediate elevations to expand their range to higher or lower elevations. Such species are most abundant at mid-elevations where contrasting environmental stresses are alleviated, contributing to the peak in diversity. The lower environmental stress at these mid-gradient sites allows for multiple phenotypes to persist, maintaining the high diversity at these sites.

### 4.3 Trends in diversity reflect changing environmental conditions

Alongside the shifts in plant function that covaried with the environmental factors along the gradient, plant community composition and diversity also change, with highest taxonomic, functional and spectral diversity at middle elevations. Mid-gradient peaks in taxonomic and functional diversity indicate that the mid-elevations supported more species and may have allowed for more niche space for species with different traits. Spectral diversity trends followed similar patterns to functional diversity. This may indicate that the leaf-level spectral diversity we measured was more influenced by spectral detection of functional traits than of phylogenetic relatedness, likely because the spectral reflectance is detecting leaf chemistry and structure (Cavender-Bares et al., 2017b; Schweiger et al., 2018).

Phylogenetic richness and species richness patterns indicated that the pool of species at higher elevations was limited to fewer lineages. Some lineages were present only at the lowest elevation, increasing the phylogenetic species richness. The significant Blomberg’s K values for RWC, LMA, LWP, and leaf nitrogen suggest that these traits are conserved. As phylogenetic richness decreased at high elevation, these traits may be especially important for survival in high alpine environments, and may be shared among close relatives.

Discrepancies arose in the direction of the patterns of diversity when abundance was accounted for. At higher elevations, the communities were mostly dominated by cushion plants that facilitate the survival of other species despite stressful abiotic conditions (Cavieres et al., 2007; Gavini, Ezcurra, & Aizen, 2020). Species evenness decreases with increasing elevation, suggesting that at higher elevations there were a few highly abundant taxa, with other rare species in the community. The difference in functional evenness and phylogenetic evenness patterns was due to uneven species abundances, driven by the presence of two highly abundant and distantly related small herb species at the 3000 m site. The mid-gradient peak in functional evenness however, suggests that traits were distributed equally throughout the community at that elevation, despite uneven species abundances. The mid-gradient peaks we observed were consistent with trends in species richness in alpine environments globally (Testolin et al., 2021). Taxonomic, phylogenetic, and functional trait composition increased with elevation, indicating that communities showed significant compositional change with increasing elevation. Though not quite statistically significant at p<0.05, the trend in spectral composition indicates that the lower elevation was different from the other, higher elevations. Considered with the environmental trends, the patterns of diversity and community composition suggested that environmental harshness may limit the plant community at the low and high end of the gradient.

### 4.4 Future directions

Climate change will affect the temperature, nitrogen availability, and water availability along the elevation gradient. We will need more research to understand the community dynamics, especially dispersal, that may affect the habitat range of species restricted to higher elevations and may change community composition along the elevation gradient. Furthermore, while we focused solely on environmental filters influencing community assembly, there are a number of other filters that impact plant communities in mountains, including plant-plant interactions and dispersal. For example, nurse plant interactions can facilitate higher levels of diversity in communities at higher elevations (Cavieres et al., 2014; Gavini et al., 2020; Pashirzad, Ejtehadi, Vaezi, & Shefferson, 2019), however competition at lower elevations can limit species from higher elevations to migrate down the gradient (Choler, Michalet, & Callaway, 2001). Dispersal limitation also influences plant communities, and accounting for seed size may help predict dispersal ability (Eriksson, 2000). We need a clear understanding of species dispersal abilities in addition to environmental filters, which will determine the available species pool for future community composition (Alexander et al., 2018). Differences in dispersal ability could influence plant diversity along elevation gradients by affecting which species can reach different elevations, while environmental filters impact which species survive.

Furthermore, trait plasticity could allow some species to respond to changing climatic conditions. Soil and plant transplant studies should be implemented in alpine regions to see the consequences of warming and reduced water on soil processes and determine the limits of plant trait plasticity.

## 5. Conclusion

We found evidence that environmental filters influence community assembly and diversity in the high alpine of the central Chilean Andes. Our results show that lower elevations were drier and higher elevations were the coldest and least nutrient rich. These environmental factors constituted harsh conditions that limited which plants could persist, based on their functional adaptations to aridity or cold. At high elevations, low nitrogen availability and temperature appear to select for species with high LMA and small leaves that can accumulate nitrogen and avert enzymatic limitation of photosynthetic rates. Intermediate elevations, where the aridity and cold stresses were alleviated, had the highest plant taxonomic, functional, and spectral diversity. Elevational patterns of trait variation in *P. secunda* mirrored those across species. Our results indicate that environmental stress and resource availability drive variation in plant function and affect which species and lineages can occur along the gradient. As the climate warms, these stresses are likely to shift upwards in elevation, affecting plant communities in different ways. By using spectral data to predict plant traits, we can rapidly determine plant community function. Understanding how environmental conditions influence plant diversity in alpine systems is an important step toward predicting how they might be affected by climate change and developing management strategies for conserving these threatened ecosystems.

## Acknowledgements

We would like to thank individuals who contributed to data collection and analysis including Megan Erding, who aided with preparation for leaf C:N analysis, Shan Kothari and Gerard Sapes for the thoughtful comments on the manuscript, and Dr. Jesus Pinto-Ledezma who advised and assisted with analysis in R. We would like to thank our funding sources, including FONDECYT project number 1180454 (MTKA), NSF DEB 1342872 (JCB) cofunded by the National Aeronautic and Space Agency through the Dimensions of Biodiversity program, NSF DBI award 2021898 (JCB) and ANID PIA APOYO CCTE AFB170008 (MTKA,PJA,VR), Grant ANID PIA/BASAL FB210006 (MTKA,PJA,VR) and Technological Centers of Excellence with Basal Financing ANID-Chile to the Cape Horn International Center (CHIC-ANID PIA/BASAL PFB210018) (MTKA,PJA,VR).

