## Supplement for "Drivers of plant diversity, community composition, functional traits and soil processes along an alpine gradient in the central Chilean Andes"

### Supplemental methods

#### Net N mineralization detailed methods

An initial extraction of field-moist soils using a 2M KCl solution was done within 4 days of soil collection for analysis of net nitrogen mineralization potential. The remainder was incubated for 30 days at 18 °C in the dark. After the incubation period, the soil was extracted in a 2M KCl solution, filtered with Whatman grade 42 filter paper and the extract frozen until analysis could be completed. Ammonium, nitrate, and nitrite concentrations were measured optically with microplate salicylate and sulfanilamide methods respectively, using a BioTek Synergy H1 microplate reader (Winooski, Vermont, USA).

#### PCR detailed methods

PCR used a final volume of 30 µl, which contained 4 µl DNA (25 ng/µl), 8.35 µl distilled water, 3 µl MgCl2 (25 mM), 6 µl buffer, 2.4 µl of dNTP (1 mM), 1.8 µl of each primer (10x), 2.4 µl BSA (25 mM) and 0.25 µl GoTaq (5 U/ µl). DNA was denatured at 95°C for 5 min, followed by 35 amplification cycles of 45 s at 94°C, annealing for 1 min at 50°C, elongation for 1.5 min at 72°C and a final extension of 7 min at 72°C. For *rbc*L, DNA was denatured at 95°C for 3 min. Followed by 30 amplification cycles of 1 min at 95°C, annealing for 45 s at 50°C, elongation for 1 min at 65°C and a final extension of 5 min at 65°C, and *matK*, DNA was denatured at 95°C for 3 min. Followed by 36 amplification cycles of 30 s at 94°C, annealing for 40 s at 58°C, elongation for 1 min at 72°C and a final extension of 10 min at 72°C.

**Bayesian analysis detailed methods**

For the combined analysis with Bayesian inference, three partitions were used corresponding for each genes, in which evolutionary models for each one were: GTR + I + G in ITS; GTR + I + G in *rbc*L; and GTR + G in *matK*. Runs appeared stationary prior to $20^6$ generations, and we conservatively excluded the first 2.0 X $10^6$ generations of each run as burn-in. Nodes with 0.95 were considered to be supported for posterior probabilities (Ronquist et al., 2012). The Tracer program (v1.6 Rambaut *et al.*, 2014) was used to visualize output parameters in order to prove stationarity and whether there are duplicated runs to converge on the same mean likelihood. The effective sample size (ESS) value was greater than 200 in a range between 406 and 17171.

**PLSR detailed methods**

The spectra were resampled to every 20 nm between 1100 and 2400 nm. We split 80% of the nitrogen data into the training group, reserving the remaining 20% for testing the model. We then ran the PLSR model using the plsr() function from the R package pls (Hovde Liland, Mevik, Wehrens, & Hiemstra, 2021) on the training group for 1000 simulations with leave one out validation and fitted with an orthogonal scores algorithm. We conducted a Tukey test on the models created with each number of components to determine how many components are needed to predict the variation in the data. We selected 11 components to use for the final model, by choosing on the lowest number of components that gave the lowest predicted residual error sum of squares (PRESS) where there was no statistical difference between the number of components. The 11 component model was validated with the testing data (see supplemental figure 3 for fit). We validated the final model by running another PLSR model including the testing data this time, with 11 components.

**<u>Supplemental figures and tables</u>**

Supplement Table 1. Taxa, herbarium voucher numbers and GenBank accession numbers.

| **Taxa** | **Voucher** | **GenBank** | | |
|---|---|---|---|---|
| | | **ITS** | *rbc*L | *matK* |
| *Acaena alpina* Poepp. ex Walp. | 184996 | MH781148 | ON542548 | ON542515 |
| *Acaena pinnatifida* Ruiz & Pav. | | MH781149 | MF963399 | MF963761 |
| *Acaena splendens* Hook. & Arn | 184995 | MH781150 | ON542549 | ON542516 |
| *Adesmia glomerula* Clos | | MH781156 | MZ198417 | |
| *Adesmia montana* Phil. | | MH781157 | MZ198416 | |
| *Adesmia schneideri* Phil | 185059 | MH781159 | ON542550 | ON542517 |
| *Alstroemeria exserens* Meyen | | MH792061 | JQ404670 | |
| *Anarthrophyllum cumingii* (Hook. & Arn.) F.Phil. | 184986 | MH781160 | ON542551 | ON542518 |
| *Azorella madreporica* Clos | 185063 | MH781165 | MZ198403 | ON542519 |
| *Azorella ruizii* G.M.Plunkett & A.N.Nicolas | 191861 | KM671870 | DQ133815 | ON542520 |
| *Berberis empetrifolia* Lam. | | | MZ198467 | |

| | | | | |
|---|---|---|---|---|
| *Calceolaria arachnoidea* Graham | 190301 | ON521128 | AY423108 | ON542521 |
| *Cerastium arvense* Cham. & Schltdl. | 184989 | | ON542552 | ON542522 |
| *Chaetanthera euphrasioides* Reiche | 191803 | DQ355866 | ON542553 | ON542523 |
| *Chenopodium philippianum* Aellen | | KF709219 | | |
| *Chuquiraga oppositifolia* D.Don | | EU841151 | EU841109 | EU841332 |
| *Collomia biflora* (Ruiz & Pav.) Brand | 185087 | MH781180 | ON542554 | HQ116935 |
| *Convolvulus demissus* Choisy | | KC528836 | KC529198 | KC529039 |
| *Ephedra chilensis* C.Presl | | | AY492036 | AY492012 |
| *Euphorbia collina* Willd. Ex Ledeb. | 190283 | | ON542555 | |
| *Galium gilliesii* Hook. & Arn. | 190326 | ON521129 | | |
| *Haplopappus anthylloides* Meyen & Walp. | 185055 | MH781202 | ON542556 | ON542524 |
| *Haplopappus schumannii* (Kuntze) G.K.Br. & W.D.Clark | 185092 | MH781204 | ON542557 | ON542525 |
| *Hordeum comosum* J. Presl | | AJ607876 | AY137441 | AB078097 |
| *Hypochaeris clarionoides* (J. Remy) Reiche | 190361 | ON521130 | | AF528408 |
| *Jaborosa caulescens* Gillies & Hook. | 185041 | MH781206 | ON542558 | ON542526 |
| *Latace andina* (Poepp.) Sassone | | KF171082 | | |
| *Loasa caespitosa* Phil. | 191792/190397 | ON521131 | ON542559 | ON542527 |
| *Melosperma andicola* Benth. | | MH781214 | MZ198432 | AY492153 |
| *Microsteris gracilis* (Hook.) Greene | 184982 | MH781217 | ON542560 | ON542528 |

| | | | | |
|---|---|---|---|---|
| *Montiopsis gilliesii* (Hook. & Arn.) D.I.Ford | | DQ090406 | | |
| *Mutisia sinuata* Cav. | | MH781221 | EU841128 | EU841355 |
| *Nassauvia looseri* Cabrera | | MG432167 | EU841132 | EU841358 |
| *Nassauvia pyramidalis* Meyen | | MG432171 | | EU841359 |
| *Nastanthus ventosus* (Meyen) Miers | 185039 | | | ON542529 |
| *Nicotiana corymbosa* J. Remy | 185007 | MH781227 | ON542561 | ON542530 |
| *Noccaea magellanica* (Pers.) Holub | 185067 | | | ON542531 |
| *Oreopolus glacialis* (Poepp. & Endl.) Ricardi | | MH095807 | | |
| *Oriastrum chilense* Wedd. | 191823 | DQ355916 | ON542562 | ON542532 |
| *Oxalis compacta* Gilles | 185002 | MH781235 | ON542563 | ON542533 |
| *Oxalis squamata* Zucc. | 185011 | MH781237 | JN587339 | ON542534 |
| *Pappostipa chrysophylla* (E.Desv.) Romasch. | | EU489113 | | EU489194 |
| *Patosia clandestina* Buchenau | | AY973514 | U49225 | |
| *Perezia carthamoides* Hook. & Arn. | 191797 | FJ979641 | EU841130 | ON542535 |
| *Phacelia secunda* J.F.Gmel. | 191801 | ON521132 | ON542564 | ON542536 |
| *Poa holciformis* J.Presl | | GQ324512 | | |
| *Polygonum bowenkampi* Phil. | 185069 | MH781241 | | ON542537 |
| *Pozoa coriacea* Lag. | 185016 | MH781242 | ON542565 | ON542538 |
| *Quinchamalium chilense* Willd. | | | EF464533 | EF464514 |
| *Quinchamalium parviflorum* Phil. | | MH781244 | | |

| | | | | |
|---|---|---|---|---|
| *Rhodolirium montanum* Phil. | 184997 | MH781245 | | ON542539 |
| *Rytidosperma violaceum* (E.Desv.) Nicora | | EU401404 | | |
| *Sanicula graveolens* Poepp. ex DC. | 191804 | ON521133 | MF963118 | ON542540 |
| *Senecio bustillosianus* J.Remy | 185056 | MH781248 | ON542566 | ON542541 |
| *Senecio pentaphyllus* Phil. | 185029 | MH781252 | ON542567 | ON542542 |
| *Sisyrinchium arenarium* Poepp. | 185052 | MH781255 | ON542568 | ON542543 |
| *Sisyrinchium cuspidatum* Poepp. | 185023 | MH781256 | ON542569 | ON542544 |
| *Stachys philippiana* Vatke | 185100 | MH781259 | ON542570 | ON542545 |
| *Taraxacum officinale* F.H.Wigg. | | KT249884 | KM361005 | MG947034 |
| *Tetraglochin alata* (Gillies ex Hook. & Arn.) Kuntze | 185051 | MH781260 | ON542571 | ON542546 |
| *Trisetum preslei* E.Desv. | | KU883528 | AY395565 | KU883578 |
| *Tropaeolum polyphyllum* Cav. | 190299 | AF254041 | AF254043 | ON542547 |
| *Viola atropurpurea* Leyb. | 185060 | MH781264 | ON542572 | |
| *Viola philippii* Leybold | 185015 | MH792062 | ON542573 | |
| *Ginkgo biloba* L. | | EF372233 | AJ235804 | EF468640 |

Supplemental table 2. Phylogenetic signal of spectral indices related to various aspects of plant physiology. Blomberg's K values compare observed signal in traits to signal predicted by a white noise (WN) null model and a Brownian motion (BM) model of evolution. K values closer to 0 correspond to random pattern of evolution, K values >1 indicate phylogenetic conservatism of traits. P value is the p value of the test for non-random signal in the phylogenetically independent contrast and the null distribution of the WN and BM models.

| Trait | K | WN p value | BM p value |
|---|---|---|---|
| RWC | 0.115479 | 0.002 | 0.000999001 |
| LMA | 0.093785 | 0.014 | 0.000999001 |
| LWP | 0.09164 | 0.015 | 0.000999001 |
| Leaf N | 0.074616 | 0.045 | 0.000999001 |
| NDMI | 0.068488 | 0.06 | 0.000999001 |
| PRI | 0.051591 | 0.213 | 0.000999001 |
| ChlNDI | 0.036536 | 0.321 | 0.000999001 |

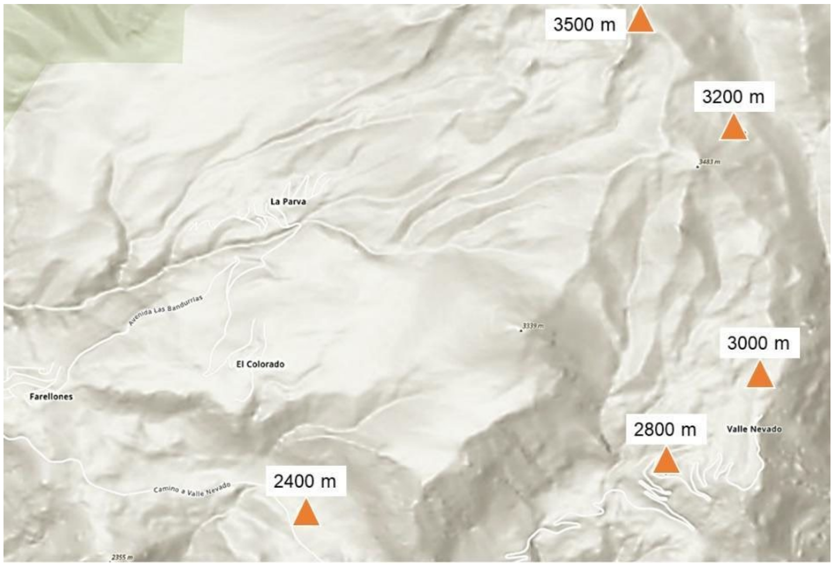


Supplemental Figure 1. Relief map of sites near Farellones used for this study.

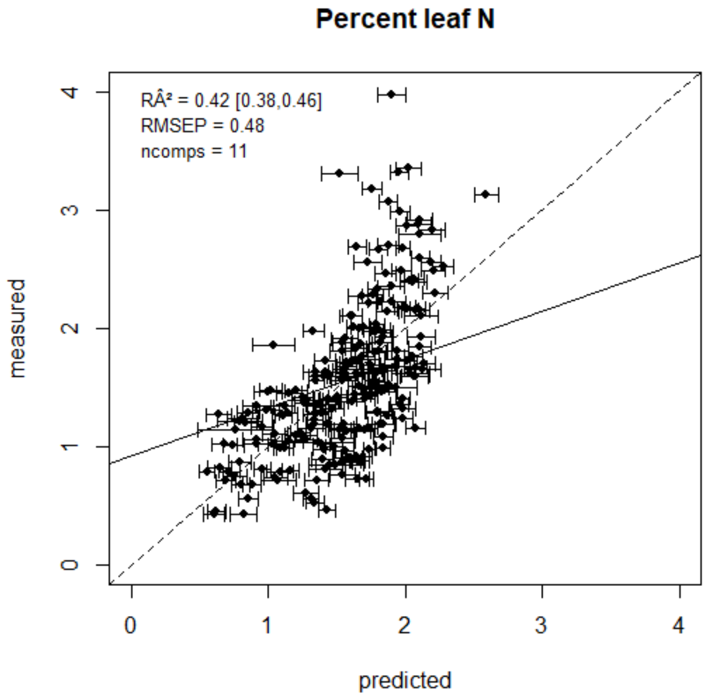


Supplemental figure 2. Predicted values of leaf N from spectra using PLSR. Dashed line indicates 1:1 fit, solid line indicates the fit of our model. The number of components (ncomps)

was determined to be 11 based on the point of plateau of PRESS values. Standardized RMSEP = 30.9 and RA2 = 0.42, with confidence intervals of 0.38-0.46.

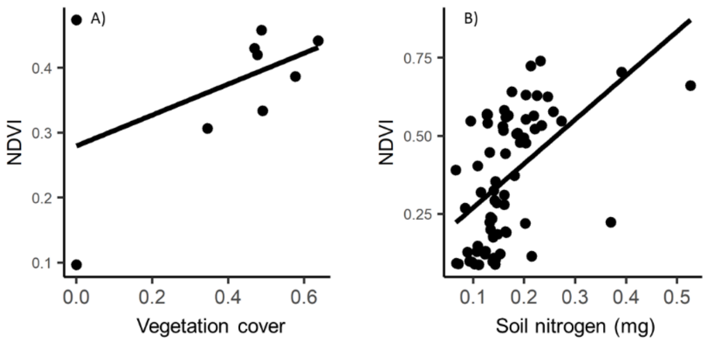


Supplemental figure 3. A) Normalized difference vegetation index (NDVI) calculated from canopy spectra on transects at 2800 m compared to percent vegetation cover. B) NDVI compared to bulk nitrogen in the soil under the vegetation.
